# Complete blood count and selected serum parameters of Venus transgenic rabbits

**DOI:** 10.1101/2022.09.14.507894

**Authors:** Bálint Biró, Gabriella Skoda, Orsolya Ivett Hoffmann, László Hiripi, Elen Gócza, Nándor Lipták

## Abstract

Green fluorescent protein (GFP) transgenic laboratory animals (mice, rats, rabbits etc.) are commonly used in basic research for modelling human diseases, studying organ development, cell transfer during pregnancy or tissue engineering. The expression of the fluorescent protein can be either ubiquitous or tissue-specific, depending on the transgenic construct and the integration site of the transgene. Despite the wide applications, the data about the physiological parameters of GFP transgenic animals are limited. In most of the transgenic lines, GFP does not induce any detrimental effect, but GFP-induced conditions are also reported. Altered T-lymphopoiesis and low white blood cell (WBC) count were observed in human ubiquitin C promoter-driven GFP transgenic (UBC-GFP) mice due to latent stem cell defect. The aim of the present study was to examine the effects of the Venus fluorescent protein on hematopoiesis and general health of transgenic rabbits, thus, hematology along with selected serum parameters were measured.

## Introduction

GFP and enhanced GFP (EGFP) transgenic mice with ubiquitous fluorophore expression were created by Okabe and his co-workers in the 1990s and this innovation made an enormous impact on biomedical research during the following decades (Ikawa *et al*. 1995; Okabe *et al*. 1997). That GFP expression pattern was provided by the chicken beta-actin promoter and cytomegalovirus enhancer (CAG) transgenic construct (Niwa *et al*. 1991). Unfortunately, homozygote EGFP transgenic mice are not viable due to position effect (Okabe *et al*. 1997), thus, researchers can use only hemizygote transgenic littermates (CAG-EGFP mice hereafter). Another transgenic line, the UBC-GFP mice also showed similar tissue distribution of the GFP, but also high levels of the fluorescence in hematopoietic cells, such as B and T lymphocytes, and dendritic cells (Schaefer *et al*. 2001). Both hemizygote and homozygote transgenic UBC-GFP mice were viable and fertile and GFP fluorescence in WBCs were stronger in the homozygotes in a genotype-dependent manner. UBC-GFP mice proved to be useful for studying of hematopoiesis, however the excessive amount of GFP in the bone marrow caused stem cell defect and led to low number of WBCs and T-lymphocytes in the peripheral blood (Faltusova *et al*. 2018; Faltusova *et al*. 2020).

In this study, CAG-Venus hemizygote and homozygote transgenic rabbits (Katter *et al*. 2013) were studied for the potential appearance of leukopenia and lymphopoiesis. Neither fecundity nor life span were affected by the transgene insertion (Liptak *et al*. 2017). Venus is a yellow shifted variant of the commonly used GFP (Shimomura *et al*. 1962) with a better resistance to acidosis and Cl^−^ (Nagai *et al*. 2002). Similarly to UBC-GFP mice, stronger fluorescence was observed by FACs analysis in peripheral blood WBCs of Venus homozygote transgenic rabbits compared with Venus hemizygote littermates (Liptak *et al*. 2017).

## Materials and methods

### Ethics statement

All experiments were approved by the Animal Care and Ethics Committee of the NARIC-Agricultural Biotechnology Institute and the Pest County’s governmental office (permission numbers: PEI/001/857-3/2015; PE/EA/610-8/2018). The experiments were complied with the Hungarian Code of Practice for the Care and Use of Animals for Scientific Purposes, including conditions for animal welfare and handling prior to slaughter.

### Blood sample collection and analysis

WT New Zealand white (n=6), Venus transgenic hemizygote (n=6) and Venus transgenic homozygote (n=5) rabbits were subjected to blood sample collection. All rabbits were male, between 3-5 months of age. Peripheral blood samples were obtained into K3-EDTA tubes (Microvette, Sarstedt, Germany, ref: 20.1341) from ear arteries of rabbits for hematology analysis (Jenkins 2008). Blood samplings were performed between 13.00-14.00 h to avoid fluctuations caused by circadian rhythm. Parameters were counted by Abacus Junior Vet5 hematology analyzer. Differential leukocyte counts were evaluated by counting 100 cells in blood smears stained with Diff-Quick.

Blood samples were also collected from the same rabbits into serum separator tubes for checking their health status (creatinine, albumin, cholesterol and triglycerides, BD Vacutainer, ref: 367953).

Both hematology and serum biochemistry analysis were done by the researchers of University of Veterinary Medicine, Department of Clinical Pathology and Oncology, Budapest, Hungary.

### Statistical analysis

Statistical analyses were performed by one way ANOVA followed by Tukey post-hoc test using PSPP Statistical Analysis Software 0.8.4. A probability value, P < 0.05 was considered statistically significant.

## Results

## Discussion

GFP transgenic animals are valuable tools in biomedical research, however, overexpression of GFP or its derivatives may contribute to the progression of certain diseases. Ubiquitous fluorophore expression evoked mild proteinuria and mild glomerulosclerosis in CAG-EGFP mice (Guo *et al*. 2007) and CAG-Venus rabbits (Lipták *et al*. 2018), while tissue-specific expression caused dilated cardiomyopathy (Huang *et al*. 2000); growth retardation (Krestel *et al*. 2004), etc. (reviewed: (Lipták *et al*. 2019)). In the majority of the studies, positive correlation between the level of GFP expression in certain tissues and the severity of diseases were observed.

In our experiments, hematology parameters of all rabbits were within reference ranges, significant differences were detected only in neutrophil counts, relative to total number of WBCs (**Table 1**). Moreover, all selected serum biochemistry parameters were also within physiological ranges, thus, rabbits had no any severe diseases (**Table 2**). Normal serum biochemistry and hematology parameters were also observed in CAG-EGFP transgenic cattle (Yum *et al*. 2018), EGFP transgenic piglets (Carter *et al*. 2002) and DsRed transgenic pigs (Chou *et al*. 2014).

**Table 1:** Complete blood count of CAG-Venus rabbits Significant difference was observed only in neutrophil counts. Asterisk represents significant difference (one way ANOVA, Tukey post hoc): * F_(2,14)_=3.83, p=0.038 compared to control bucks. Data are presented as mean ± standard deviation (SD). The neutrophil mean values of all groups were within normal reference ranges (neutrophil: 20-75% of total WBCs).

| **Hematologic parameters** | **WT control (n=6)** | **CAG-Venus hemizygote (n=6)** | **CAG-Venus homozygote (n=5)** |
| --- | --- | --- | --- |
| **RBC ($10^{12}/L$)** | 5.92 ± 1.29 | 6.34 ± 1.48 | 5.95 ± 0.30 |
| **Hgb (g/L)** | 110.67 ± 24.31 | 118.60 ± 22.86 | 114.80 ± 5.26 |
| **Ht (L/L)** | 0.37 ± 0.09 | 0.40 ± 0.07 | 0.37 ± 0.02 |
| **MCV (fL)** | 62.83 ± 3.43 | 64.33 ± 4.97 | 62.20 ± 1.64 |
| **MCH (pg)** | 18.73 ± 2.05 | 19.12 ± 2.45 | 19.36 ± 1.29 |
| **MCHC (g/L)** | 299.00 ± 28.14 | 296.67 ± 22.54 | 310.40 ± 15.04 |
| **PLT ($10^9/l$)** | 269.14 ± 136.5 | 336.83 ± 136.2 | 212.20 ± 86.47 |
| **WBC ($10^9/l$)** | 8.61 ± 3.90 | 8.53 ± 2.60 | 8.02 ± 0.77 |
| **Neutrophil (%)** | 36.83 ± 10.70 | 56.17 ± 13.79* | 46.20 ± 11.50 |
| **Lymphocyte (%)** | 59.17 ± 12.22 | 41.50 ± 12.80 | 52.40 ± 12.18 |
| **Monocyte (%)** | 2.33 ± 2.83 | 2.33 ± 2.73 | 1.20 ± 1.10 |
| **Eosinophil (%)** | 0.40 ± 0.89 | 0.00 ± 0.00 | 0.40 ± 0.89 |

Abbreviations: red blood cell (RBC), hemoglobin (Hb), hematocrit (Hct), mean cellular volume (MCV), mean cellular hemoglobin (MCH), mean cellular hemoglobin concentration (MCHC), platelet (PLT), white blood cell (WBC),

**Table 2:** Serum biochemistry of CAG-Venus rabbits Significant differences were not observed between any groups in any parameters. Data are presented as mean ± SD. The values of all rabbits were within normal reference ranges (Melillo 2007).

| **Serum biochemistry parameters** | **WT control (n=6)** | **CAG-Venus hemizygote (n=6)** | **CAG-Venus homozygote (n=5)** |
| --- | --- | --- | --- |
| **Albumin (g/l)** | 40.85 ± 4.66 | 42.35 ± 3.00 | 42.68 ± 1.90 |
| **Creatinine (umol/l)** | 110.83 ± 22.94 | 103.83 ± 22.94 | 113.00 ± 22.00 |
| **Cholesterol (mmol/l)** | 1.32 ± 0.37 | 1.52 ± 0.46 | 1.36 ± 0.43 |
| **Triglycerides (mmol/l)** | 0.98 ± 0.62 | 1.07 ± 0.59 | 0.65 ± 0.07 |

In a previous report, UBC-GFP mice successfully engrafted the transplanted bone marrow and spleen of WT mice without any conditioning (Faltusova *et al*. 2018). Significantly lower number of WBCs and T-lymphocytes were observed in peripheral blood of UBC-GFP mice by the same research group. This phenomena could be caused by the overexpression of GFP *per se* in the bone marrow stem cells (Faltusova *et al*. 2020). The UBC-GFP transgene was integrated into a noncoding region on mouse chromosome 17, but the position effect, as a reason of defect in lymphopoiesis cannot be ruled out completely (Liu *et al*. 2020; Necas *et al*. 2021). Excessive Venus fluorescence was not observed in the bone marrow of CAG-Venus rabbits (data not shown). In CAG-EGFP mice, total body X-ray irradiation led to increased apoptosis in the bone marrow cells, compared with WT mice (Liu *et al*. 2020). For the present study, the potential differences in bone marrow apoptosis between WT and CAG-Venus rabbits after X-ray irradiation could not be examined due to the 3R regulations. The increased reactive oxygen species (ROS) production in the bone marrow of CAG-EGFP mice could be responsible for the aforementioned increased cell death, according to a recent report (Liu *et al*. 2021). During the maturation of EGFP, H_2_O_2_ is produced by the transgenic cell, in 1:1 ratio (Zhang *et al*. 2006).

Besides defective polyubiquitination, oxidative stress in GFP-expressing cells could be the main reason for the reported pathological events in transgenic animals (Baens *et al*. 2006; Ganini *et al*. 2017).

In conclusion, altered hematopoiesis was not observed in Venus transgenic rabbits. Hematology and serum biochemistry parameters should be measured in newly created GFP transgenic lines and the experiments should be designed cautiously if any alteration is detected.

## Acknowledgements

This work was supported by National Research, Development and Innovation Office (NKFIH) grant no. 108921, NVKP_16-1-2016-0039 and 120870.

